# Repositioning the Sm-binding site in *S. cerevisiae* telomerase RNA reveals RNP organizational flexibility and Sm-directed 3′-end formation

**DOI:** 10.1101/167361

**Authors:** Evan P. Hass, David C. Zappulla

## Abstract

Telomerase RNA contains a template for synthesizing telomeric DNA by reverse transcription and has been proposed to act as a flexible scaffold for holoenzyme protein subunits in the RNP. In *Saccharomyces cerevisiae*, the telomerase subunits Est1 and Ku bind to the telomerase RNA, TLC1, and it has been shown that these proteins still function when their binding sites are repositioned within the RNA. TLC1 is also bound by the Sm_7_ protein complex, which is required for stabilization of the predominant, non-polyadenylated (poly(A)–) TLC1 isoform. Here, we first show that Sm_7_ can perform this function even when its binding site is repositioned via circular permutation to several different positions within TLC1, further supporting the conclusion that the telomerase holoenzyme is organizationally flexible. Next, we tested the hypothesis that the location of the Sm_7_-binding site relative to the 3′ end is contrastingly important. When we moved the Sm site to locations 5′ of its native position, we observed that this stabilized shorter forms of poly(A)– TLC1 in a manner precisely corresponding to how far upstream the Sm site was moved. This provides strong evidence that the location of Sm_7_ binding to TLC1 controls where the mature poly(A)– 3′ end is formed. In summary, our results show that Sm_7_ and the 3′ end of yeast telomerase RNA comprise an organizationally flexible module within the telomerase RNP and provide insights into the mechanistic role of Sm_7_ in telomerase RNA biogenesis.

## INTRODUCTION

Telomeres are regions of repetitive sequence at the ends of eukaryotic chromosomes that buffer against shortening caused by the end-replication problem. In most eukaryotes, telomeres are lengthened by telomerase, an RNP enzyme that adds new telomeric repeats to the ends of telomeres via reverse transcription of an RNA template (Greider and Blackburn 1985). Fundamentally, this process is carried out by the two core subunits of telomerase: telomerase reverse transcriptase (TERT) and the non-coding telomerase RNA. TERT is the catalytic protein subunit of telomerase, while the telomerase RNA contains the template for reverse transcription of telomeric repeats (Shippen-Lentz and Blackburn 1990) as well as binding sites for accessory protein subunits of the telomerase RNP (Seto et al. 1999; Peterson et al. 2001; Seto et al. 2002; Lemieux et al. 2016).

In *Saccharomyces cerevisiae*, the major isoform of telomerase RNA (TLC1) is 1157 nucleotides long and is predicted to form a Y-shaped secondary structure, with the template and TERT-binding regions in the central core and binding sites for holoenzyme subunits towards the tips of each arm (Dandjinou et al. 2004; Zappulla and Cech 2004) (Fig. 1B). Like all telomerase RNAs, yeast telomerase RNAs are evolving very rapidly in sequence and size, especially in the regions between protein-binding sites (Tzfati et al. 2000; Zappulla and Cech 2004). These intervening regions can be deleted in TLC1, resulting in miniaturized “Mini-T” RNAs that still function *in vivo* despite being as little as one-third the size of wild-type TLC1 (Zappulla et al. 2005). It has also been shown that two important holoenzyme subunits, Est1 and Ku, retain function when their binding sites in TLC1 RNA are repositioned (Zappulla and Cech 2004; Zappulla et al. 2011). These findings show that the telomerase RNP exhibits a high degree of organizational flexibility, and they have led to the model that TLC1 acts as a “flexible scaffold” — the RNA brings together the protein subunits to form the holoenzyme but does not need to hold them in a specific position relative to one another or the catalytic core (Zappulla and Cech 2004; Zappulla and Cech 2006; Zappulla et al. 2011; Lebo and Zappulla 2012).

In addition to Est1, Ku, and TERT (Est2), TLC1 is also bound by the Sm_7_ complex (Seto et al. 1999). The Sm_7_ heteroheptameric protein complex is involved in biogenesis and stabilization of most spliceosomal snRNAs (Jones and Guthrie 1990; Will and Luhrmann 2001). Sm_7_ binds to the consensus sequence AU_5-6_GR (Branlant et al. 1982; Liautard et al. 1982; Mattaj and De Robertis 1985; Hamm et al. 1987), which is present in TLC1 at nucleotides 1143–1150 (Fig. 1A) (Seto et al. 1999). This site is located just 7 nucleotides 5′ of position 1157, which is the 3′ end of poly(A)–TLC1 (Bosoy et al. 2003), the aforementioned most-abundant (“major”) isoform of TLC1. A less-abundant (“minor”) isoform, poly(A)+ TLC1, contains an extra ~100 nucleotides of TLC1 sequence on its 3′ end, as well as a poly(A) tail (Chapon et al. 1997). In addition, there are very low-abundance TLC1 transcripts terminated by the Nrd1-Nab3-Sen1 (NNS) complex that only have ~50 extra nucleotides beyond the poly(A)–TLC1 3′ end at position 1157 (Jamonnak et al. 2011; Noel et al. 2012), but these transcripts are presumably not stable and are not detectable by northern blot.

**Figure 1.**
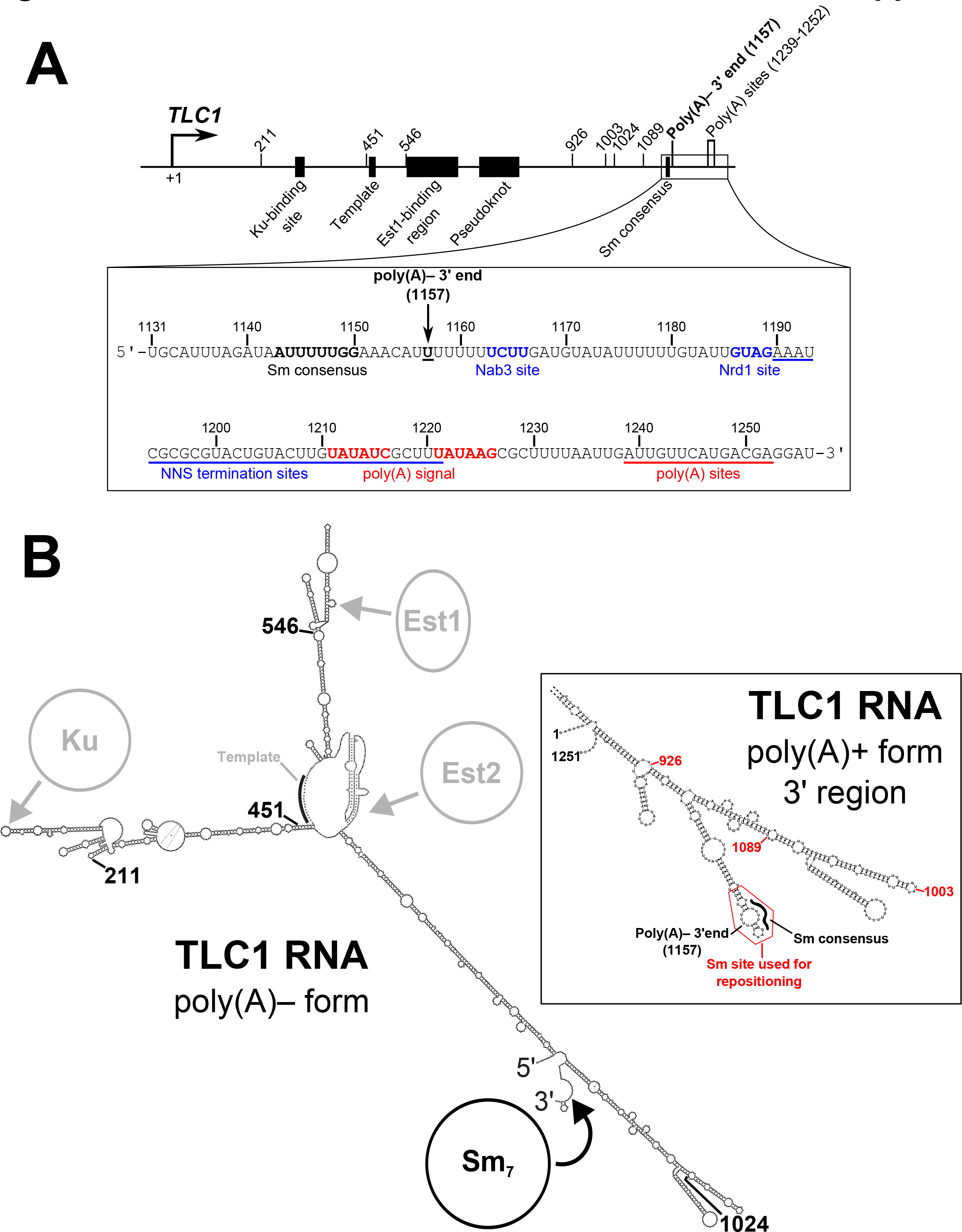
TLC1 sequence schematic and secondary-structure models. (A) Schematic of the *TLC1* gene sequence. The *TLC1* regions encoding the Ku-binding site, template, Est1-binding region, pseudoknot, and Sm consensus in the RNA are denoted as black rectangles. Locations of all Sm-repositioning sites are noted by tick marks, as are the poly(A)– 3′ end and the polyadenylation sites. The inset shows the RNA sequence of the TLC1 3′ region. The Sm consensus and poly(A)– 3′ end are noted in bold. The Nab3 and Nrd1 binding sites are bolded in blue, and the region containing NNS termination sites is underlined in blue (Jamonnak et al. 2011; Noel et al. 2012). Similarly, the polyadenylation signal is red boldface, and the region containing polyadenylation are underscored by a red line (Chapon et al. 1997). (B) Secondary structure models of 1157- and 1251-nt forms of TLC1. The pictured secondary structure of poly(A)– TLC1 is based on previously published models (Dandjinou et al. 2004; Zappulla and Cech 2004; Niederer and Zappulla 2015). The location of Sm binding is indicated in black as are the four locations to which the Sm binding site and the 5′ and 3′ ends are repositioned in the SmCP alleles used in Figure 2. The binding regions for Ku, Est1, and Est2 and the template region are indicated in gray. The inset shows the secondary structure model of the 3′ region of poly(A)+ TLC1 based on a previously published model (Zappulla and Cech 2004). The portion of the RNA used when repositioning the Sm-binding site (nucleotides 1138 to 1165) is outlined in red. The locations of the Sm consensus and poly(A)– 3′ end are noted in black, and the three positions to which this Sm-binding site was repositioned in Figure 3 are indicated in red.

In wild-type cells, the major, poly(A)–TLC1 isoform is present at ~29 molecules per cell, while poly(A)+ TLC1 is at only ~1 molecule per cell (Mozdy and Cech 2006). However, when the Sm consensus in TLC1 is mutated, only the minor poly(A)+ TLC1 isoform is detectable, likely because poly(A)– TLC1 is not stable without Sm_7_ bound (Seto et al. 1999). Due to this critically low abundance of telomerase RNA, these mutant cells (*tlc1-Sm^−^* cells) display senescence-related growth defects (e.g., small or mixed colony sizes) but do not display a fully senescent phenotype (Seto et al. 1999). This “near-senescent” growth phenotype seems to be in agreement with the fact that poly(A)+ TLC1 abundance averages only ~1 molecule per cell. Because a single molecule of TLC1 per cell is the average over a population of cells, some cells in the population probably have fewer molecules than average (i.e., none) and will eventually senesce, while others cells in the same population will have more than one molecule of TLC1 and will be able to lengthen their telomeres enough to continue dividing.

Although it was shown 18 years ago that Sm_7_ binds to telomerase RNA in yeast (Seto et al. 1999), several questions about Sm function in the telomerase RNP remain unanswered. First, while it has been shown that Sm_7_ can retain function when repositioned within Mini-T along with repositioning of the 5′ and 3′ ends of the RNA (also called “circular permutation”) (Mefford et al. 2013), the flexible scaffold model has not been tested for Sm_7_ in full-length TLC1 as it has been for Est1 and Ku. Additionally, there are several open mechanistic questions regarding Sm_7_ function in TLC1 biogenesis. It was proposed that TLC1 is initially transcribed as poly(A)+ TLC1 and that this RNA is then processed into poly(A)– TLC1 (Chapon et al. 1997). It has also been proposed that the nuclear exosome exonucleolytically trims poly(A)+ TLC1 from 3′ to 5′ and is then sterically blocked at nucleotide 1157 by Sm_7_, thus generating poly(A)– TLC1 (Coy et al. 2013). Although it has been shown that nuclear exosome mutants accumulate more poly(A)+ TLC1 than wild type (Coy et al. 2013), the hypothesis that Sm_7_ defines the mature 3′ end of poly(A)– TLC1 via this mechanism has remained untested. It also remains unclear whether poly(A)+ TLC1 is in fact the precursor of poly(A)–TLC1 or if NNS-terminated TLC1 transcripts are processed into the poly(A)– isoform.

Here, we show that Sm_7_ retains its function in telomerase when repositioned via circular permutation in full-length TLC1, demonstrating that the Sm complex and the RNA ends are an organizationally flexible module on the telomerase RNP’s RNA scaffold. Having shown that the Sm-binding site can function at diverse positions within circularly permuted TLC1 alleles, we next used Sm-site repositioning in the context of the unpermuted RNA to test the hypothesis that the Sm binding position defines the mature 3′ end of poly(A)–TLC1. When we repositioned the Sm site further 5′ in telomerase RNA, the stabilized poly(A)– TLC1 RNAs were correspondingly shorter than wild-type poly(A)– TLC1. This shows that Sm_7_, in addition to providing stability, dictates formation of the mature end of this RNA just 3′ of its binding site.

## RESULTS

### Sm_7_ retains function when its binding site is repositioned by circular permutation in TLC1

To test if the Sm_7_ protein complex retains its functions in the telomerase RNP when its binding site is repositioned in TLC1, we chose to reposition the Sm-binding site and the 3′ end together by circular permutation. Thus, <u>Sm</u> repositioning by <u>c</u>ircular <u>p</u>ermutation (“SmCP”) alleles allow assessment of Sm functions at new locations in the RNP while retaining Sm binding site location relative to the 3′ end of the RNA. We designed these *TLC1-SmCP* alleles using the TLC1 RNA secondary structure as a guide. As shown in Figure 2A, position 1134 was fused to position 1, thus excising the endogenous Sm site from its native location along with the downstream transcriptional termination sequences (Fig. 1A). Then, newly encoded ends were introduced at 4 different positions in the *TLC1* sequence, while the Sm-binding region and transcriptional termination sequences (nucleotide 1130 to the end of the TLC1 locus) were appended to the new 3′ end of the gene. Additionally, to maintain endogenous expression of TLC1, the *TLC1* promoter and first 10 nucleotides from the wild-type 5′ end were retained at the 5′ end of the new gene. SmCP alleles were created in this manner at positions 211, 451, and 1024 (Figs. 1, 2A) — the same three positions used to reposition the Est1-binding region previously (Zappulla and Cech 2004). Furthermore, in order to test the positional flexibility of Sm function in all three arms of TLC1, we also created an SmCP allele in the Est1 arm of TLC1, at position 546.

**Figure 2.**
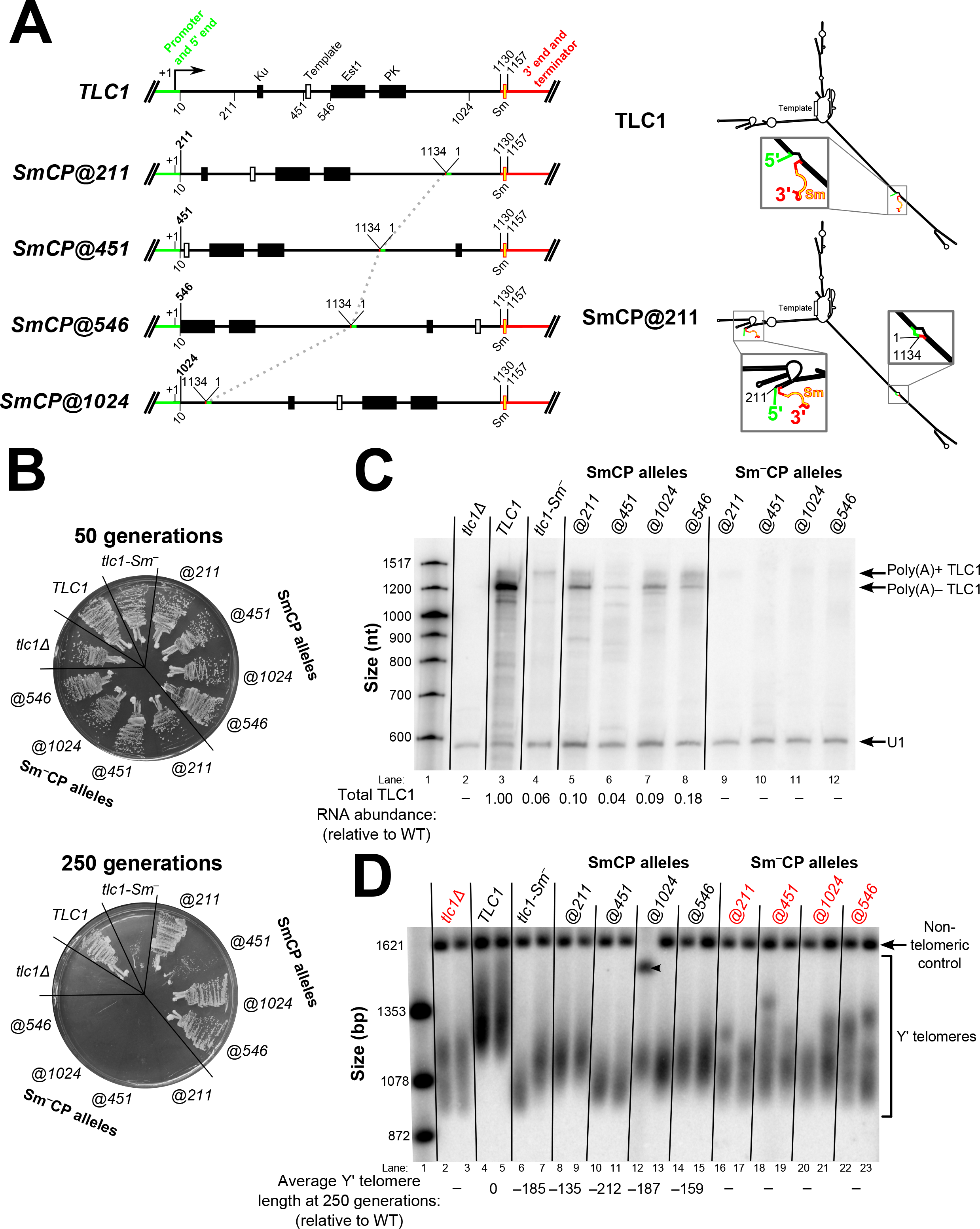
Sm_7_ retains function in telomerase RNA when its binding site is repositioned via circular permutation. (A) Schematics of *SmCP* gene construction and expected RNA structure. The native ends of the *TLC1* gene were effectively sealed off by connecting nucleotide 1134 with nucleotide 1, thus removing the Sm site from its native position in the RNA. A dashed line connects the location of this 1134-to-1 fusion between the 4 circularly permuted alleles. The wild-type *TLC1* promoter through nucleotide 10 (green) as well as the sequence from nucleotide 1130 on (red) flank the circularly permutated central 1–1130 region, thereby repositioning the Sm-binding site (yellow) along with the 5′ and 3′ ends in the encoded transcript. Black rectangles indicate the Ku-binding site, Est1-binding region, and pseudoknot (PK). A white rectangle indicates the template. The secondary-structure models of wild-type TLC1 and an example of an SmCP (TLC1-SmCP@211) are schematized to the right, using the same coloring scheme as in the gene diagrams to the left. Details of the secondary structure of these large RNAs are omitted in these low-resolution schematics for the sake of simplicity, but, in fact, these regions of TLC1 have wild-type sequence and predicted secondary structure very similar to wild type as well. (B) *TLC1-SmCP* alleles with a wild-type Sm-binding site support sustained growth and do not cause cells to senesce. All *TLC1* alleles were expressed from centromeric plasmids in a *tlc1*Δ *rad52*Δ background, and cells were serially passaged on solid media for ~250 generations (10 re-streaks). (C) Sm_7_ confers poly(A)–telomerase RNA stabilization and appropriate 3′-end formation when repositioned via circular permutation. Total RNA was isolated from cells used in the passaging experiment shown in Figure 2B and analyzed by northern blotting with TLC1 and U1 snRNA probes. Total TLC1 RNA abundance was normalized to U1 abundance to control for sample loading in each lane and then set relative to the wild-type condition. The numbers displayed below the blot are averages of two independent biological replicates. (D) SmCP alleles support stable, short telomeres. Genomic DNA was isolated from cells in the passaging experiment in Figure 2B at ~250 generations for non-senescent conditions (labeled in black) and ~75 generations for senescent conditions (labeled in red). Telomere length was then analyzed by Southern blotting. The blot was probed for telomeric sequence and for a 1621-bp non-telomeric restriction fragment (non-telomeric control). The pairs of lanes in the blot shown are independent biological-replicate samples, and the telomere length numbers are averages of the two replicates except in the *TLC1-SmCP@1024* condition. In this condition, the telomere length could not be quantified in the first replicate due to anomalous migration of the non-telomeric control (black arrowhead), so the displayed telomere length number in this condition is a quantitation of only the second replicate sample.

First, to test if these SmCP alleles maintain telomerase function and prevent senescence like wild-type *TLC1*, these RNAs were expressed from the *TLC1* promoter on a centromeric plasmid in a *tlc1*Δ background, and growth was monitored for~250 generations. Notably, *TLC1-SmCP@211, @546*, and *@1024* all supported wild-type growth throughout the ~250 cell divisions (Fig. 2B). This shows that the Sm_7_ complex is functioning when its binding site is moved to these three different locations in TLC1. In contrast to these three alleles, one Sm site-repositioning circular permutant, *TLC1-SmCP@451*, led to a near-senescent phenotype very similar to *tlc1-Sm^−^* cells, causing mixed or small colony sizes after ~125 generations. However, when compared with the growth phenotype observed when the Sm site was mutated in this allele, this result suggests that Sm_7_ in fact retains partial function in the SmCP@451 telomerase RNP. To control for whether the SmCP alleles affected growth for reasons other than Sm function, we also created “Sm^−^CP” alleles, in which the repositioned Sm-binding consensus was mutated so that it is rendered binding-incompetent (Raker et al. 1999; Seto et al. 1999). Unlike the near-senescent cells expressing the conventional *tlc1-Sm*^−^ allele, and in contrast to the sustained viability of cells expressing the SmCP alleles, cells expressing any of the 4 Sm^−^CP alleles completely senesced by ~150 generations. The long-term viability of all SmCP strains compared to the senescence of all Sm^−^CP strains shows that the Sm_7_ protein complex retains its functions when bound at each of the four different positions in the circularly permuted TLC1 RNAs. Furthermore, these are the first circular permutants of full-length TLC1 RNA to be tested, and it is noteworthy that the telomerase RNA ends themselves can be repositioned to any of these four locations while permitting telomerase functionality *in vivo*. However, because unpermuted TLC1 with a mutated Sm site (tlc1-Sm^−^) does not quite cause a senescent phenotype, the contrasting senescent phenotype of all of the Sm^−^CP alleles suggests that circular permutation of the RNA interferes at least slightly with TLC1 function and/or accumulation.

Next, we assessed how TLC1 RNA processing and abundance was affected in the SmCP alleles. For all four SmCP alleles, the poly(A)– TLC1 isoform — predicted to be just 15 nt longer than wild type — was readily detectable by northern blotting, although at a lower abundance than wild type (Fig. 2C). This shows that Sm_7_ can perform its function in TLC1 3′-end processing and (to a lesser extent) RNA accumulation when its binding site is repositioned. The particularly low abundance of the SmCP@451 poly(A)– RNA probably contributes to why these cells exhibit a near-senescent phenotype. When we assessed TLC1 RNA abundance in the four Sm^−^CP alleles, we observed that poly(A)+ TLC1 was essentially undetectable, unlike in cells containing the un-permuted *tlc1-Sm*^−^ allele (Fig. 2C; compare lanes 9–12 with lane 4). This shows that while cells expressing tlc1-Sm^−^ accumulate just enough functional TLC1 RNA (poly(A)+) to prevent complete senescence, circularly permuting TLC1 without also including a functional Sm-binding site at the new 3′ end reduces RNA abundance to negligible levels, resulting in senescence.

We next assessed the effect of the Sm site repositioning and circular permutation on telomere length. We isolated genomic DNA from *TLC1-SmCP* cells at the end of passaging (i.e., ~250 and ~75 generations for the non-senescent and senescent conditions, respectively) and subjected the DNA to Southern blotting with a telomeric probe. The results show that cells expressing *SmCP@211* and *@546* alleles had the longest telomeres, although all four SmCP constructs supported lengths substantially shorter than wild type (Fig. 2D, lanes 8–15). Overall, the telomere-length phenotypes were consistent with the cell-growth results, and they confirmed that telomeres were stably maintained through 250 generations, unlike in Sm^−^CP cells. Telomeres were longer in *TLC1-SmCP@211* and *@546* cells than in the near-senescent *tlc1-Sm*^−^ cells, providing additional evidence that Sm_7_ is functioning when repositioned. As expected, the near-senescent *TLC1-SmCP@451* cells had the shortest telomeres of any of the SmCP alleles, averaging 212 bp shorter than wild type (lanes 10 and 11). Considering that the *SmCP@451* allele supported the shortest telomeres and the lowest RNA abundance, while the *SmCP@211* and *@546* cells had the longest telomeres and the highest RNA levels, these data suggest that relocated Sm_7_ functioned best at positions 211 and 546 and that SmCP telomerase RNAs do not support full-length telomeres primarily, or potentially entirely, because of their low levels of accumulation.

### Sm binding defines the mature 3′ end of poly(A)–TLC1 RNA

It has been hypothesized that Sm_7_ binding to TLC1 nucleotides 1143–1150 defines the mature 3′ end of poly(A)–TLC1 at nucleotide 1157 by blocking exonucleolytic trimming at the terminus of an initially longer transcript (Seto et al. 1999; Coy et al. 2013). Although genetic data suggest that the nuclear exosome is involved in the maturation of poly(A)– TLC1 (Coy et al. 2013), the model that Sm-binding defines the mature 3′ end of poly(A)– TLC1 has not been rigorously evaluated. Having observed that Sm_7_ functions when its binding site is repositioned in the telomerase RNA, we next tested the hypothesis that Sm_7_ controls 3′-end formation in poly(A)– TLC1 by repositioning the Sm-binding site further 5′ in TLC1 without circular permutation. If the Sm-binding site’s position defines the location of the 3′ end of poly(A)–TLC1, repositioning the Sm-binding site to a position further 5′ in the RNA should correspondingly result in a truncated poly(A)–form. In the published secondary-structure model of poly(A)+ TLC1 (nts 1–1251), the Sm-binding consensus is on the 5′ portion of an internal loop on the side of a hairpin, while nucleotide 1157 (the poly(A)– 3′ end) is predicted to be on the other side of this loop, directly across from the Sm-binding consensus (Fig. 1B, inset) (Zappulla and Cech 2004). Additionally, the nucleotides in and around this putative internal loop show higher sequence conservation (Zappulla and Cech 2004; Mefford et al. 2013), suggesting that this predicted structure as a whole, not just the 8-nucleotide Sm-binding consensus, is required for Sm_7_ to perform its function. Thus, based on this conservation and predicted local secondary structure, we chose nucleotides 1138 to 1165 as the RNA module that would most likely be functional when repositioned within TLC1. We then inserted this Sm-binding site into tlc1-Sm^−^ at positions 1089, 1003, and 926 (Fig. 1B, inset). All three of these positions are predicted to be within bulged loops located on the distal half of the terminal arm, a region that is dispensable for telomerase function (Zappulla et al. 2005).

Cells expressing TLC1 RNAs with these relocated Sm sites were passaged for 250 generations to assay for senescence. The strains expressing tlc1-Sm^−^+Sm@1089 or @926 displayed wild-type growth, whereas *tlc1-Sm^−^+Sm@1003* cells exhibited a near-senescent growth phenotype, similar to *tlc1-Sm^−^* cells (Fig. 3A). Northern blotting analysis revealed that, in agreement with the observed growth phenotypes, the Sm-binding sites inserted at positions 1089 and 926 in tlc1-Sm^−^ were functional in stabilizing poly(A)–TLC1 (at 1% and 7% of wild-type poly(A)–TLC1 abundance, respectively), while the Sm-binding site inserted at position 1003 was not (Fig. 3B). Additionally, the poly(A)– RNAs that were stabilized by the inserted Sm sites at positions 1089 and 926 were shorter than wild-type poly(A)– TLC1 RNA (indicated by red arrowheads in lanes 8, 9, 12, and 13), and their lengths measured from the northern were within 0.1% and 1.9%, respectively, of expected sizes based on the positions of the inserted Sm sites. This result provides strong evidence that the position of Sm binding on TLC1 RNA defines the mature 3′ end of poly(A)–TLC1.

**Figure 3.**
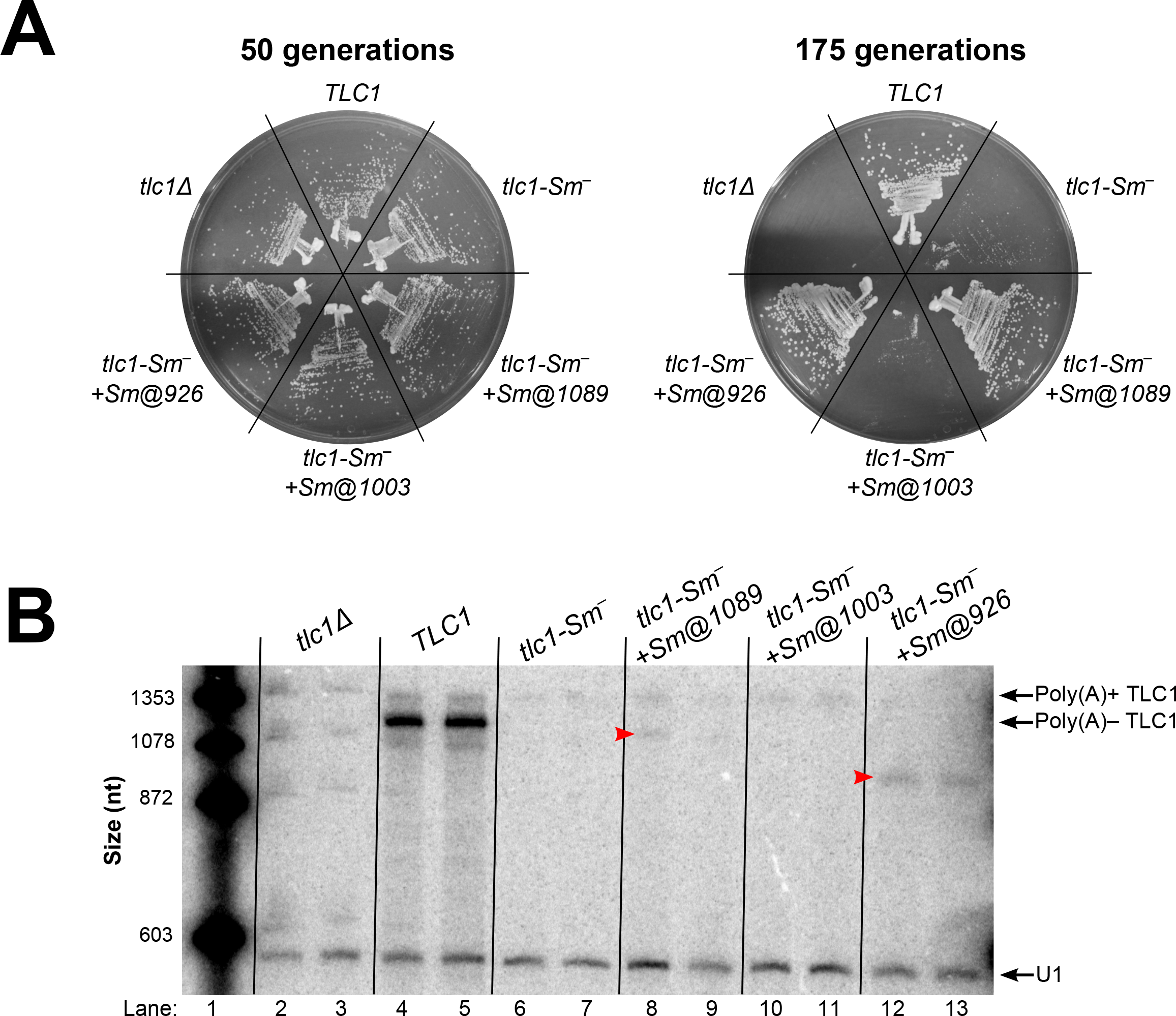
Repositioning the Sm binding site at two of three positions further 5′ in TLC1 results in stabilization of shorter poly(A)–TLC1 RNAs. (A) Sm sites inserted in tlc1-Sm^−^ restored robust growth at positions 1089 and 926, but not at 1003. *TLC1* alleles were expressed and cells were passaged as in Figure 2. (B) Functional Sm sites inserted 5′ of the native position in tlc1-Sm^−^ stabilized shorter poly(A)–TLC1 RNAs. Total RNA was isolated from cells used in the passaging experiment in Figure 3A and analyzed by northern blotting as in Figure 2C. Red arrowheads indicate the shorter poly(A)–TLC1 RNAs stabilized in *tlc1-Sm^−^+Sm@1089* and *tlc1-Sm^−^+Sm@926* cells. The pairs of lanes represent independent biological-replicate samples.

## DISCUSSION

The telomerase RNA differs in many ways from other large, well-studied, non-coding RNAs such as ribosomal and spliceosomal RNAs. Although its existence is highly conserved among eukaryotes, its sequence and length vary greatly between species (Chen and Greider 2004). Considering the rapid evolution of telomerase RNA along with experimental results has led to the model that telomerase RNA functions as a flexible scaffold for protein subunits in the telomerase RNP (Zappulla and Cech 2004; Zappulla and Cech 2006; Lebo and Zappulla 2012; Mefford et al. 2013). Thus, unlike other large RNP enzymes that require a precise structural organization of components in the complex for function (e.g., the ribosome), the *S. cerevisiae* telomerase RNP has organization that has been shown to be strikingly flexible and this has provided a paradigm for many long noncoding RNAs (Zappulla and Cech 2004; Zappulla et al. 2011; Mefford et al. 2013).

Here, we have shown that organizational flexibility of the yeast telomerase RNP extends to include the Sm_7_ subunit, which binds just before the 3′ end of its RNA subunit’s most abundant isoform and stabilizes it. Despite repositioning of the Sm site within TLC1 via circular permutation to four dramatically different locations across all three RNA arms, Sm_7_ was still able to promote processing and stabilization of the major (poly(A)–) TLC1 isoform. Although poly(A)– TLC1 RNA abundance was reduced in these SmCP alleles, Sm_7_ still retained at least partial function at all positions tested. This demonstrates that Sm_7_ and its binding site in TLC1 function as an organizationally flexible module, along with the RNA ends in the telomerase RNP. Between the results presented here and previous repositioning studies for the binding sites of Est1 and Ku subunits (Zappulla and Cech 2004; Zappulla et al. 2011), it is now evident that TLC1 is an organizationally flexible scaffold for these three well-established RNA-bound holoenzyme subunits of telomerase. While organizational flexibility has not yet been tested for the Pop1/Pop6/Pop7 complex that was recently reported to bind to an essential sequence in TLC1 near the Est1-binding site (Lemieux et al. 2016), the entire Est1-Pop1/6/7-binding region of TLC1 has been shown to function when it is expressed *in trans* as a separate RNA while Est1 is artificially tethered near the 3′ end of TLC1 (Lebo et al. 2015), suggesting that the flexible scaffold model applies to the Pop1/6/7 proteins as well.

After determining that the Sm_7_ subunit can function at several different non-native positions within the telomerase RNP in the context of circular permutation, we next used Sm-site repositioning to investigate a mechanistic hypothesis regarding Sm function in TLC1 RNA biogenesis. We tested whether the position of Sm binding defines the mature 3′ end of poly(A)–TLC1 by repositioning the Sm-binding site further 5′ in the RNA. At positions where the Sm site was functional, this resulted in stabilization of correspondingly shorter poly(A)– TLC1 RNAs, showing that Sm_7_ does indeed direct 3′-end formation of poly(A)–TLC1.

Although our experiments show clearly that Sm_7_ helps define the mature 3′ end of poly(A)–TLC1, it remains unclear what the precursor of this RNA is. While it has been proposed that poly(A)+ TLC1 is the precursor of poly(A)– TLC1 (Chapon et al. 1997; Coy et al. 2013), it is also possible that TLC1 transcripts terminated by the NNS complex are unprocessed precursors of poly(A)–TLC1 (Jamonnak et al. 2011; Noel et al. 2012). However, the Sm-binding site in these NNS-terminated TLC1 transcripts, which are 20-50 nucleotides shorter than poly(A)+ TLC1 (Jamonnak et al. 2011), likely adopts a conformation different from that in the published secondary structure model of the 1251-nt poly(A)+ TLC1 (Zappulla and Cech 2004). The most common site of NNS-mediated transcription termination in TLC1 is nucleotide 1195 (Jamonnak et al. 2011), and we observe that in an *Mfold* secondary-structure prediction of TLC1 RNA terminated at 1195, the Sm-binding consensus indeed does not form the same hairpin and internal-loop structure as that shown in the published secondary-structure model for poly(A)+ TLC1 (Fig. S1). In this secondary-structure prediction for the 1195-nt form of TLC1, the Sm-binding consensus is largely single-stranded (except for a two-base pair hairpin that seems unlikely to be stable enough to actually form), with the remaining sequence 3′ of the consensus forming a hairpin. Unlike the published secondary-structure model for poly(A)+ TLC1, this structural prediction of the 1195-nt form of TLC1 fits with the observation that many Sm-binding sites are single-stranded and are followed and/or preceded by hairpin structures (Branlant et al. 1982; Liautard et al. 1982). Given the similarities between this structural prediction and other more well-characterized Sm-binding sites, it is possible that the Sm-binding site in TLC1 adopts this conformation in at least some, if not all, unprocessed forms of TLC1.

In summary, repositioning the Sm-binding site and the ends of yeast telomerase RNA has yielded insights about the organizationally flexible nature of the RNA and the mechanistic role of Sm_7_ in its biogenesis. Sm_7_ can functionally tolerate repositioning of its binding site within TLC1 along with the RNA ends. Furthermore, the telomerase enzyme’s fundamental functions can also endure the substantial reorganization of the RNA induced by circular permutations. Sm-site repositioning also has revealed that the mature 3′ end of poly(A)– TLC1 appears to be controlled by the location of Sm_7_ binding to the RNA. The experimental approach of repositioning the Sm binding site in order to move the mature 3′ end of poly(A)–TLC1 may prove useful in identifying the precursor of poly(A)–TLC1 and fully characterizing the pathway for telomerase RNA biogenesis in budding yeasts.

## METHODS

### Construction of TLC1-SmCP alleles

The *TLC1-SmCP* alleles were constructed using a plasmid containing two tandem copies of *TLC1* (pDZ573, see Table S1 for full list of plasmids used). In this construct, the two copies of *TLC1* are fused into a single gene such that nucleotide 1134 of the first *TLC1* copy is followed directly by nucleotide 1 (see Fig. 2A) of the second *TLC1* copy, meaning that the Sm site and transcriptional termination sequences are absent in the first *TLC1* sequence. Circular permutation was performed by PCR-amplifying specific segments of this template DNA beginning at the site of repositioning in the first *TLC1* copy and ending at the same site in the second copy (e.g., for *SmCP@211*, this resulted in a PCR product containing base pairs 211–1134 followed by 1–210). These circularly permuted DNAs were inserted into a separate plasmid containing the natural ends of the *TLC1* gene such that sequences from nucleotide 10 upstream and from nucleotide 1130 downstream (which includes the Sm site) are retained at the ends of the new gene. As an example, the sequence of *TLC1-SmCP@211* is laid out from promoter to terminator as follows: the *TLC1* promoter through nucleotide 10, 211–1134, 1–210, and 1130 through the transcriptional terminator and natural end of the *TLC1* gene (see Fig. 2A). As a result of this construction scheme, nucleotides 1–10 and 1130–1134 are contained twice in all *TLC1-SmCP* alleles, making the RNAs 15 nucleotides longer than wild-type *TLC1*.

### Experiments in yeast

All experiments were performed in the strain TCy43 (*MATa ura3-53 lys2-801 ade2-101 trp1-1 his3-Δ200 leu2-Δ1 VR::ADE2-TEL adh4::URA3-TEL tlc1::LEU2 rad52::HIS3 pTLC1-LYS2-CEN*) (Seto et al. 1999). All *TLC1* alleles were expressed from centromeric plasmids (Table S1) that were derived from pSD107 (*pTLC1-TRP1-CEN)* (Diede and Gottschling 1999). TCy43 was transformed with *TLC1*-containing plasmids, and colonies were streaked to minimal –TRP–LYS medium. Loss of the *pTLC1-LYS2-CEN* cover plasmid was selected for by re-streaking cells to minimal –TRP medium containing α-aminoadipate. These cells were then serially re-streaked nine times to minimal –TRP medium and photographed after each round of growth. When estimating the number of generations at different points throughout passaging, each round of growth after loss of the cover plasmid (including the round of growth in the presence of α-aminoadipate) is approximated as 25 generations.

### Northern blotting

Northern blotting was performed as described previously (Zappulla et al. 2005; Zappulla et al. 2011; Hass and Zappulla 2015; Lebo et al. 2015). Briefly, cells from the serial passaging plates were grown in liquid cultures to an OD600 of ~1.0 and harvested. Total RNA was isolated using the hot-phenol method (Kohrer and Domdey 1991). 15–30 μg of total RNA from each sample was boiled, separated by urea-PAGE, and then transferred to Hybond-N^+^ Nylon membrane (GE). The membrane was UV-crosslinked and probed for both TLC1 and U1 snRNA sequence using 100-fold fewer counts of U1 probe than TLC1 probe to account for the large difference in abundance between the two RNAs. TLC1 RNA abundance was calculated by normalizing to U1 abundance, and numbers in the text and figures are expressed relative to the wild-type *TLC1* condition. Lengths of poly(A)–TLC1 RNAs in Fig. 3B were calculated using the molecular weight standard shown.

### Southern blotting

Southern blotting was performed as described previously (Zappulla et al. 2005; Zappulla et al. 2011; Hass and Zappulla 2015; Lebo et al. 2015). Briefly, cells were grown and harvested in the same manner as those used for northern blots, and genomic DNA was isolated from these pellets (Gentra Puregene system). Roughly equal amounts of genomic DNA were digested with XhoI and then separated on a 1.1% agarose gel. DNA was transferred to Hybond-N^+^ Nylon membrane (GE) to which it was UV-crosslinked. The membrane was probed for yeast telomeric sequence and a 1621-bp non-telomeric XhoI fragment that served as a non-telomeric control band (Friedman and Cech 1999). Average Y′ telomere length was calculated using the weighted average mobility method described previously (Zappulla et al. 2011).

## ACKNOWLEDGEMENTS

We thank members of the Zappulla laboratory for their input on these experiments, particularly Benjamin Ford and Timothy Kistner who helped with designing and cloning *TLC1* alleles used in Figure 3. This research was supported in part by the National Institute of General Medical Sciences of the National Institutes of Health under award number R01GM118757 to D.C.Z. as well as startup funds from Johns Hopkins University.

---

**Figure S1.**
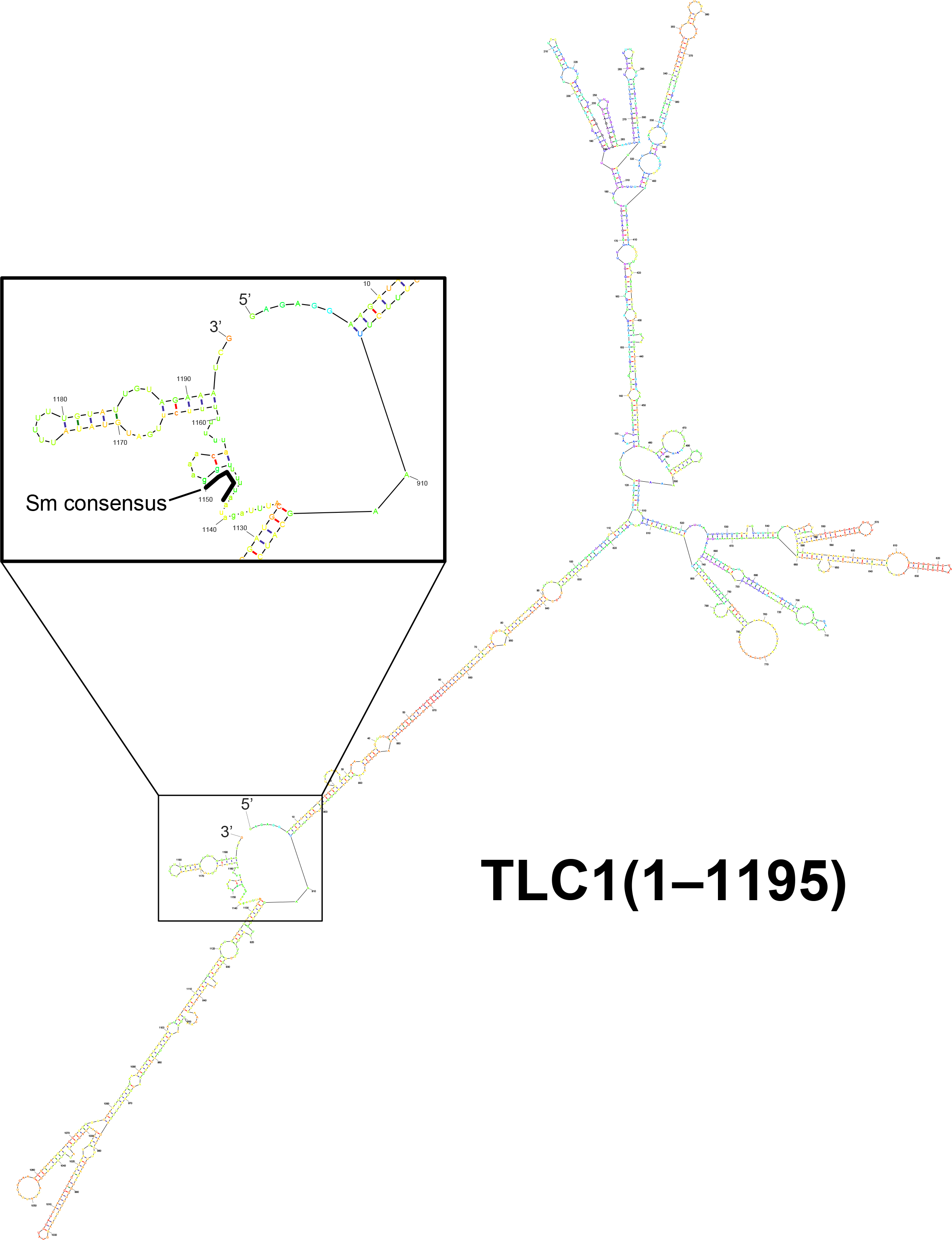
*Mfold* software prediction for the secondary structure of NNS-terminated TLC1 (nucleotides 1–1195). The structure is colored using the pnum preset on the *Mfold* RNA web server to reflect the determinedness based on free-energy calculations (Zuker and Jacobson 1998). The 3′ end of the RNA structure is enlarged in the inset, and the Sm-binding consensus is outlined in black.

